# Discovery and binding mode of small molecule inhibitors of the apo form of human TDO2

**DOI:** 10.1101/2024.01.09.574827

**Authors:** Carina Lotz-Jenne, Roland Lange, Sylvaine Cren, Geoffroy Bourquin, Laksmei Goglia, Thierry Kimmerlin, Micha Wicki, Manon Müller, Nadia Artico, Sabine Ackerknecht, Philippe Pfaff, Christoph Joesch, Aengus Mac Sweeney

## Abstract

Tryptophan-2,3-dioxygenase (TDO2) and indoleamine-2,3-dioxygenase (IDO1) catalyze the conversion of L-tryptophan to N-formyl-kynurenine and play important roles in metabolism, inflammation, and tumor immune surveillance. Their enzymatic activities depend on their heme contents, which vary dynamically according to biological conditions. Inhibitors binding to heme-containing holo-TDO2 are known, but to date no inhibitor that binds to the heme-free state (apo-TDO2) has been reported. We describe the discovery of the first apo-TDO2 targeting inhibitors, to our knowledge, together with their co-crystal structures and inhibition of cellular TDO2 activity at low nanomolar concentrations.

## Introduction

Tryptophan-2,3-dioxygenase (TDO2) and indoleamine-2,3-dioxygenase 1 (IDO1) are structurally distinct heme-binding enzymes that have evolved independently to catalyze the same reaction: the conversion of L-tryptophan (Trp) to N-formyl-kynurenine (NFK). Together, they play important roles in metabolism, inflammation, and tumor immune surveillance. Their cellular enzymatic activity levels depend on the relative proportion of apo (inactive, heme-free) and holo (active, heme containing) forms, which vary dynamically according to biological settings including nitric oxide (NO) levels [1]. Since the enzymatic activity of IDO1 and TDO2 have been shown to contribute to the immune-suppressive condition in the tumor microenvironment (reviewed in [2]), small molecule inhibitors are regarded as a therapeutic option to reinvigorate anti-tumor immunity. Small molecule, active-site inhibitors binding to apo-IDO1 [3, 4], holo-IDO1 [5–7] and holo-TDO2 [8] have been reported, as well as dual holo-IDO1/holo-TDO2 inhibitors [9–11].

Several IDO1 selective inhibitors have been tested in oncology clinical trials, often in combination with immune checkpoint inhibitors, but have failed to demonstrate sufficient efficacy to date [12]. Concerning the few dual holo-IDO1/TDO2 inhibitors taken into phase I clinical trials, results were only reported for M4112 (Merck KGaA, Darmstadt, Germany) which was discontinued due to insufficient pharmacodynamic effect [13]. Most of the clinical studies in the IDO1 field have been performed with the holo-IDO1 inhibitor epacadostat, which was also the only IDO1 inhibitor for which a phase 3 study was completed. In this clinical trial, epacadostat failed to show improved progression-free survival or overall survival compared with placebo plus the immune checkpoint inhibitor pembrolizumab in patients with unresectable or metastatic melanoma [14]. While many hypotheses have been raised as to why the trial was not successful, it appears evident by now that the dose used was insufficient to block the IDO1 enzyme [15].

Compared to holo-IDO1 inhibitors, apo-IDO1 binding inhibitors, in particular BMS-986205 (linrodostat), show higher potency in cell-based assays and *in vivo* pharmacodynamic studies [15]. In addition, apo-binders compete with heme instead of the Trp substrate and could lead to sustained IDO1 inhibition due to delayed rebinding of heme to the apoenzyme following inhibitor dissociation [3, 16]. Despite the high likelihood that the same principle and potential therapeutic benefits apply to TDO2, which also exists in both apo and holo forms, no inhibitors targeting apo-TDO2 have been reported to date. Therefore, we have screened a diverse compound library and identified the first reported inhibitors of apo-TDO2 to our knowledge. The discovery and characterization of these small molecules paves the way for a new class of TDO2 inhibitors with a different mechanism of action and the potential for greater *in vivo* efficacy.

## Results

### Identification of the inhibitors Rac-1 and Cpd-3 in a cellular TDO2 activity assay

The weak inhibitor Rac-1 (IUPAC name *rac*-3-chloro-*N*-(1-(6-chloropyridin-3-yl)-2-phenylethyl)aniline) was identified as a hit in a screening campaign of a diverse set of compounds in a human TDO2 cellular assay utilizing SW48 cells, in which endogenous TDO2 is highly expressed [8]. TDO2 expression in SW48 results in the conversion of Trp to NFK by TDO2 and the subsequent processing of NFK to kynurenine (KYN) by NFK formamidase. The potent inhibitor Cpd-4 (IUPAC name ethyl 9-(3-(2-isopropylphenyl)-1-((*S*)-1-phenylethyl)ureido)-2-methoxy-4-oxo-6,7,8,9-tetrahydro-4*H*-pyrido[1,2-*a*]pyrimidine-3-carboxylate) is a close analog of the hit Cpd-3 identified in the same screening campaign.

In this assay, which measures the accumulation of NFK and KYN in the supernatant, the racemate Rac-1 showed 60% inhibition at 10 µM. In a second experiment, in which the compound was tested in dose-response, the IC_50_ was determined at 7.5 µM, with 82% inhibition at 52 µM, the highest inhibitor concentration tested (S1 Fig 1) while the second hit (Cpd-3) and its potent analog Cpd-4 displayed IC_50_ values of 327 nM and 14.8 nM, respectively. The inhibitory effect of Rac-1, Cpd-3 and Cpd-4 was not a result of toxicity as for all three compounds the Tox IC_50_ was determined as >10 µM. While Rac-1 did not inhibit the activity of purified holo-TDO2 at concentrations up to 10 µM in a standard 30 min enzymatic assay at room temperature with 30 min pre-incubation of the compound with TDO2, Cpd-3 displayed an IC_50_ of 17.4 µM (Cpd-4 was not tested).

For the enzymatic assay, recombinant human TDO2 comprising amino acids 19-407 with a N-terminal hexahistidine tag, expressed in *Escherichia coli* and purified to homogeneity, was incubated in assay buffer consisting of 75 mM phosphate buffer at pH 7.0 supplemented with 100 µM ascorbic acid, 50 U/ml catalase, 0.01% bovine serum albumin (BSA), and 0.01% Tween 20 (protocol modified from [17]). We speculate that the absent activity of Rac-1 and the much weaker activity of the second hit in the enzymatic assay in comparison to the cellular assay may be due to the kinetics of heme dissociation from TDO2, the absence of alternative heme binding proteins in the assay system, and/or the presence of a high concentration (200 µM) of Trp substrate which likely stabilizes the holo form of TDO2.

In total, five apo-TDO2 inhibitors are discussed: the weak (micromolar) screening hit Rac-1, which is a racemic mixture; Cpd-1 and Cpd-2, the separated enantiomers of Rac-1; the nanomolar screening hit Cpd-3 and its more potent analog Cpd-4.

### Binding of Rac-1 to apo-TDO2 in living cells

To confirm binding of Rac-1 to apo-TDO2 in intact cells, we used bioluminescence resonance energy transfer (BRET) utilizing MCF7 cells expressing TDO2 genetically fused to NanoLuc luciferase and a cell-permeable fluorescent tracer derived from a compound specifically binding to the apo form of TDO2 (structure not disclosed) and inhibiting TDO2 in SW48 cells with an IC_50_ of 37 nM (S1 Fig 2). In brief, MCF7 cells, which do not express TDO2, were transduced with a lentiviral vector consisting of NanoLuc (BRET donor) fused to the C-terminus of TDO2. For the BRET tracer, we applied NanoBRET 590SE as BRET acceptor [18]. Addition of the tracer to MCF7-TDO2-NanoLuc cells under equilibrium conditions resulted in a tracer concentration dependent increase in BRET signal (Fig 1a). Adding the tracer at 6 nM and 30 nM, corresponding to EC_50_ and EC_80_, respectively, and Rac-1 in excess (10 µM) to the cells resulted in attenuation of the BRET signal (Fig 1b), confirming the ability of Rac-1 to compete with the tracer for binding to the apo form of TDO2 in intact cells. The dependence of the NanoBRET technology on a NanoLuc fusion protein as a BRET donor ensures a highly specific readout: only a BRET acceptor binding to the TDO2-NanoLuc fusion protein can produce a BRET signal.

**Fig 1.**
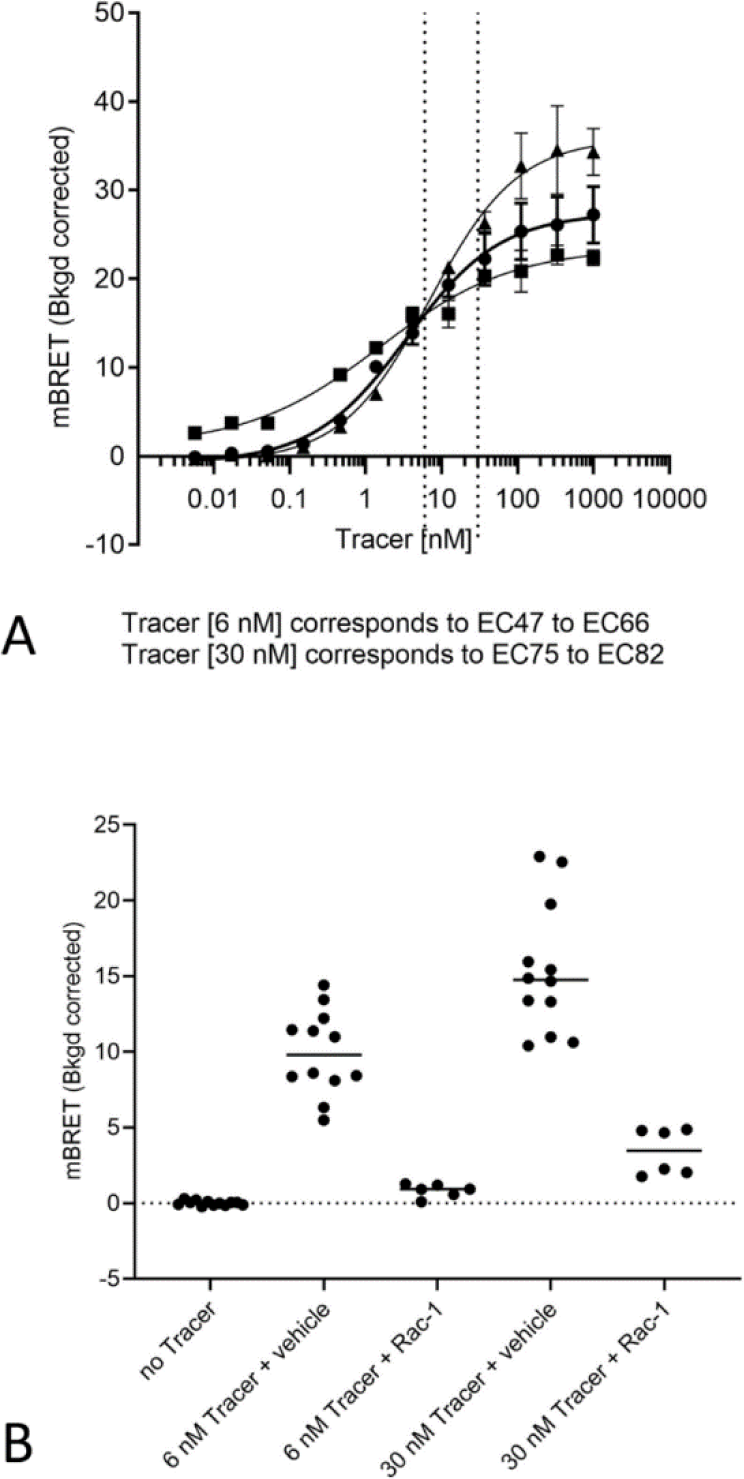
Measuring TDO2 target engagement in living cells. (A) Incubation of MCF7-TDO2-NLuc cells under equilibrium conditions with increasing concentrations of tracer resulted in increasing BRET signal. BRET values at each tracer concentration were background-corrected by subtraction of parallel measurements made in the absence of compound (vehicle) as described in Materials and Methods. Apparent tracer affinities were estimated using GraphPad “Find ECanything” curve fitting. EC corresponding to the 6 nM and 30 nM tracer concentrations used in competition with apo-TDO2 inhibitors in (B) are indicated with dotted lines. (B) Incubation of MCF7-TDO2-NLuc cells under equilibrium conditions with either 6 nM or 30 nM of tracer and 10 µM of Rac-1 results in attenuation of the BRET signal. Data from 3 independent experiments are shown in (A) and (B).

### Differential scanning fluorimetry (DSF)

We used differential scanning fluorimetry (DSF) to confirm thermal stabilization of apo-TDO2 by compound Rac-1, its separated enantiomers Cpd-1 and Cpd-2, or the more potent Cpd-3 or Cpd-4, all in the presence of the known exosite binding compound α-methyltryptophan (AMT). An exosite that binds both the substrate Trp and its α-methyl analog AMT has been identified previously in holo-TDO2 [19]. Biochemical and cellular analyses indicated that Trp binding to the exosite of holo-TDO2, while not affecting the catalytic properties, prevented its proteolytic degradation by the ubiquitin-dependent proteasomal pathway. Trp binds with 100-fold higher affinity at the holo-TDO2 exosite than at the active site, with approximate K_D_ values of 0.5 µM and 54 µM, respectively [19]. AMT binding stabilized the enzyme against heat, urea and proteases [19].

DSF measurements demonstrated that AMT also stabilizes the apo form of TDO2 (Table 1). We observed high initial fluorescence values (below the apo-TDO2 melting temperature) in the absence of an apo-TDO2 inhibitor, which we speculate is due to binding of the Sypro orange dye in the unoccupied hydrophobic heme binding pocket. AMT led to an improved apo-TDO2 melting curve quality and was therefore included in all apo-TDO2 inhibitor DSF measurements. The inhibitor Rac-1 showed stabilization of two different apo-TDO2 constructs in DSF (Table 1). Cpd-1, tentatively assigned as the S-enantiomer, showed greater stabilization of apo-TDO2 (Table 1) in agreement with the clear electron density for this compound in the co-crystal structure. Cpd-2, the R-enantiomer, showed weaker stabilization of apo-TDO2 (Table 1) and was not observed in co-crystal structures. By far the most pronounced stabilization of apo-TDO2 (ΔT_m_ of 14.5 °C) was caused by the addition of the potent inhibitor Cpd-3.

**Table 1.** Stabilization of two different TDO2 constructs (short, aa 39-389; long, aa 19-406) by the addition of Rac-1 or Cpds 1 to 4. The mean value of n replicates is shown, with n shown in the right column. The Tm of TDO2 in the presence of 500 µM AMT (shown in bold) was used as the reference value for ΔTm calculation, resulting in a negative ΔTm for the only sample without AMT.

| Protein | AMT (μM) | Compound | Conc (μM) | T<sub>m</sub> ($^{\circ}$C) | ΔT<sub>m</sub> ($^{\circ}$C) | n |
| --- | --- | --- | --- | --- | --- | --- |
| TDO2 short | 0 | - | - | 50.08 | -1.76 | 6 |
| **TDO2 short** | 500 | - | - | 51.84 | - | 6 |
| TDO2 short | 500 | **Rac-1** | 400 | 57.68 | 4.08 | 3 |
| TDO2 short | 500 | **Cpd-1** | 400 | 60.20 | 6.60 | 3 |
| TDO2 short | 500 | **Cpd-2** | 400 | 54.67 | 2.83 | 3 |
| **TDO2 long** | 500 | - | - | 58.37 | - |  |
| TDO2 long | 500 | **Rac-1** | 50 | 61.52 | 3.29 | 2 |
| TDO2 long | 500 | **Rac-1** | 200 | 62.80 | 4.31 | 2 |
| **TDO2 long** | 500 | - | - | 59.17 | - |  |
| TDO2 long | 500 | **Cpd-3** | 25 | 70.43 | 11.26 | 2 |
| TDO2 long | 500 | **Cpd-4** | 25 | 73.65 | 14.48 | 2 |

### Monitoring heme-competition by holo-TDO2 UV-VIS absorbance

To explore whether the identified apo-binders can compete with heme-binding to TDO2, a UV-VIS absorption spectrum of holo-TDO2 was measured in the absence or presence of compounds. It is well known that heme-containing proteins exhibit a characteristic absorbance peak at 405 nm, the so called Soret band, which corresponds to the electronic state of the iron. A shifted Soret peak indicates a protein-ligand interaction, and a reduced intensity can reveal heme loss. While the holo-TDO2 absorption spectrum in the presence of the potent Cpd-4 showed a clear time-dependent reduction in the Soret band intensity similar to what has been reported for known IDO apo-binders [16], no reduction was seen in the presence of Rac-1 or the vehicle control (Fig 2).

**Fig. 2.**
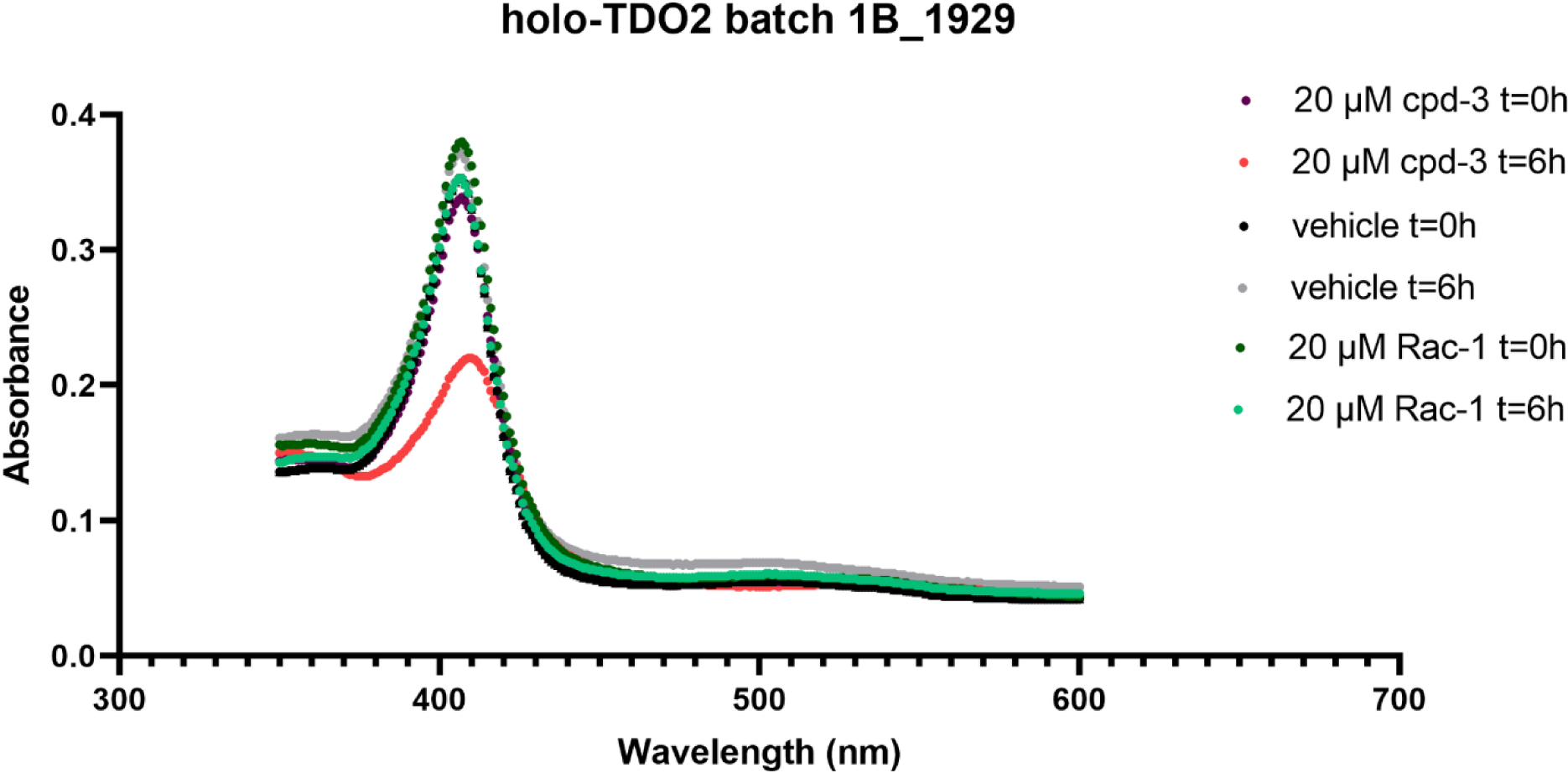
The UV-VIS absorption spectrum of holo-TDO2 in the presence of Cpd-4, Rac-1 or the vehicle control. The incubation of 5 µM holo-TDO2 with 20 µM Cpd-4 shows a reduction of the absorbance peak at 405 nm (Soret band) after 6 h of coincubation at 37 °C indicating heme loss due to compound binding (red dotted line). No reduction of the Soret band was seen with the vehicle control (grey dotted line) or 20 µM Rac-1 (green dotted line).

The weak potency of Rac-1 and high holo-TDO2 concentration used in the assay most likely explain this finding.

### Structure of the inhibitor Rac-1 bound to apo-TDO

Apo-TDO2 was expressed using an *E. coli* strain that is unable to synthesize heme. In the presence of AMT, which was included due to the stabilizing effect observed in DSF experiments, apo-TDO2 could be crystallized alone and in the presence of the active site inhibitor Rac-1, and later with the purified single enantiomers Cpd-1 and Cpd-2. The structure of the racemic mixture Rac-1 in complex with apo-TDO2 confirmed the presence of the inhibitor in the heme binding pocket. Visual inspection of the electron density indicated that the S-enantiomer was more likely to be the predominantly bound enantiomer, supported by a marginally higher real space correlation coefficient (RSCC) for the S-enantiomer (best fit 0.958, mean of four molecules 0.953) than for the R-enantiomer (best fit 0.950, mean of 4 molecules 0.947) after refinement. Chiral separation of the enantiomers and DSF measurements confirmed the preferential binding of one enantiomer (Table 1). Cpd-1, as the purified enantiomer which showed greater thermal stabilization of TDO2, was therefore tentatively assigned as the S-enantiomer based on the inhibitor electron density (S1 Fig 3). The lower resolution co-crystal structure of this purified enantiomer confirmed the binding mode (S1 Fig 4) but could not be used to assign the stereochemistry unequivocally, while the less stabilizing enantiomer (Cpd-2, tentatively assigned as the R-enantiomer) was not observed in co-crystal structures. As the resolution of the Rac-1 structure was higher than that of Cpd-1 (2.62 Å vs 3.05 Å), the Rac-1 structure has been used for analysis and figures.

The 2.62 Å resolution structure of Rac-1 revealed that the compound binds in the heme binding pocket of apo-TDO2 (Fig 3A). The inhibitor is bound to all four TDO2 molecules of the homo-tetramer. The overall conformation of apo-TDO2 in the inhibitor complex resembles that of the 2.90 Å structure of apo-TDO2 alone (PDB ID 4PW8) [20], but with a significant inhibitor-induced rearrangement of the flexible loop 148-156 at the heme binding site (Fig 3B). This first apo-TDO2 structure was published several years before the discovery of apo-IDO1 inhibitors, and before the importance of the apo forms of IDO1 and TDO2 *in vivo* was understood, therefore the potential relevance of the observed apo form of TDO2 was not discussed.

**Fig 3.**
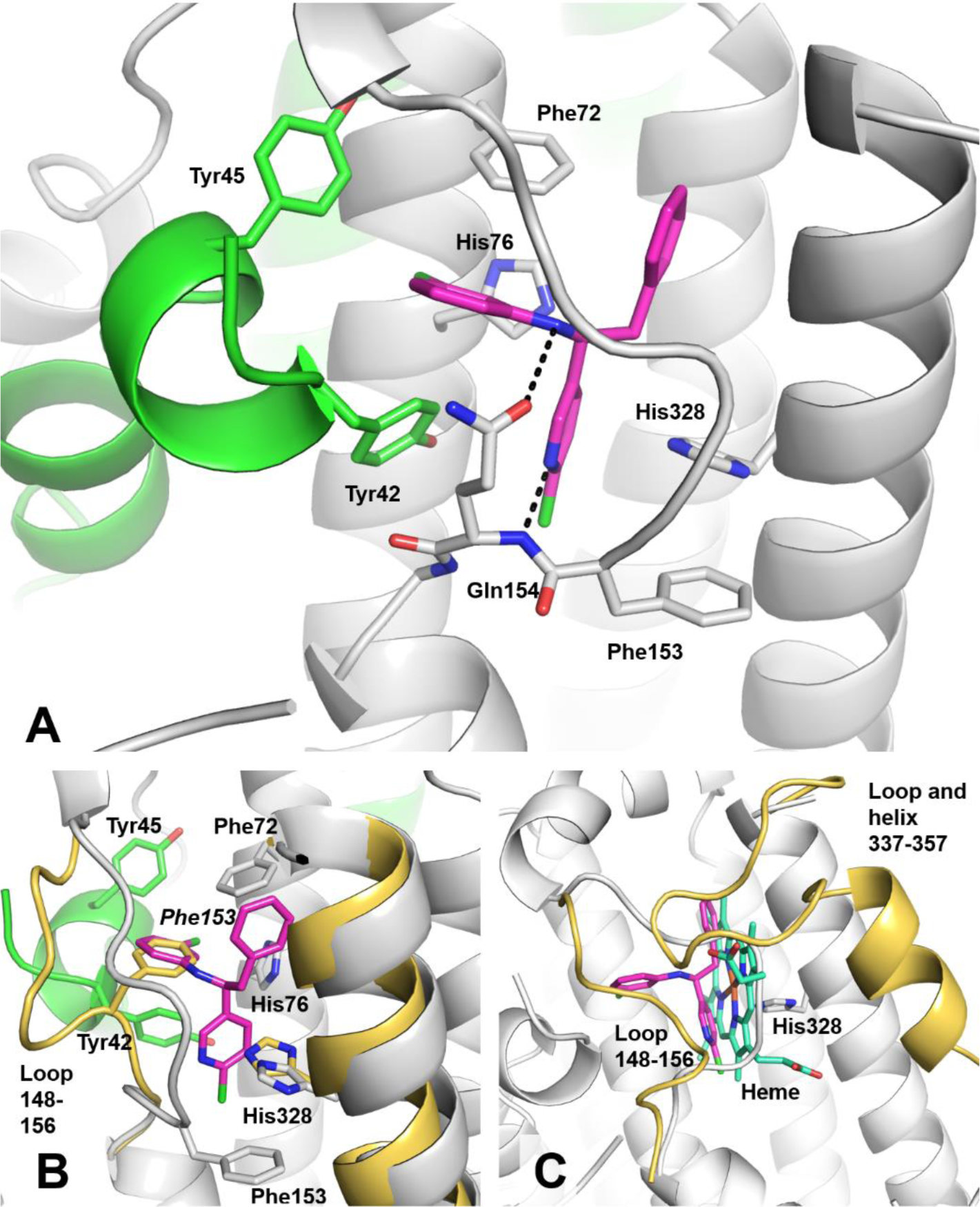
(A) Binding interactions of the inhibitor (magenta) in the heme binding pocket of apo-TDO. The binding pocket of TDO2 is formed by two TDO2 monomers, shown in grey and green cartoon. Hydrogen bond interactions between the inhibitor and Gln154 are shown as black dashes. (B) A superposition of the inhibitor bound apo-TDO2 structure (protein in grey and green, inhibitor in magenta) with the published structure of apo-TDO2 (4PW8, shown in yellow), showing the conformational rearrangement of loop 148-156. Phe153 of the 4PW8 structure (labeled in italic) is replaced by the closely matching chlorobenzyl of the inhibitor. (C) A superposition of the inhibitor bound apo-TDO2 structure (both protein chains in grey for clarity, inhibitor in magenta) with the published structure of tryptophan bound holo-TDO2 (5TIA, heme shown in cyan and residues 148-156 and 337-357 shown in yellow). In stick format, non-carbon atoms are colored blue (N), red (O) and green (Cl).

The inhibitor binding site is formed by two TDO2 molecules, and TDO2 molecule A is shown in green in Fig 3. In a conformational rearrangement most likely driven by hydrogen bond interactions between the inhibitor and the main chain (3.0 Å) and side chain (3.5 Å) of Gln154 (Fig 3A), loop 148-156 closes tightly around the inhibitor. The binding pocket contains many hydrophobic residues in the direct vicinity of the inhibitor, including four phenylalanines (45, 72, 140 and 158), two leucines (46 and 132), two methionines (331 and 335) and a tryptophan (324). The highly hydrophobic nature of the inhibitor binding pocket is likely to contribute significantly to inhibitor binding.

A superposition of the inhibitor complex with the published apo-TDO2 structure [20] shows that Phe153 is displaced by the inhibitor, with a C_α_ shift of 6 Å. The resulting nascent pocket is filled by the chlorobenzyl group of the inhibitor, which binds analogously to the (now displaced) benzyl ring of Phe153 (Fig 3B). The side chain of His328 is displaced to a lesser extent by the inhibitor chloropyridine, with C_α_ and side chain shifts of approximately 1 and 3 Å, respectively. His238 displacement is associated with a slight outward shift of helix residues 325-333 (Fig 3B), to an intermediate position lying approximately halfway between the apo-TDO2 (4PW8) and holo-TDO2 structures (such as PDB entry ID 5TIA) [19].

The Rac-1 inhibitor complex also differs significantly from the structure of holo-TDO2 (Fig 3C) in the presence or absence of bound holo-TDO2 inhibitors [19]. The flexible loop 148-156 of holo-TDO2 adopts a very open conformation to accommodate the bulky heme, in contrast to the tightly closed conformation induced by Rac-1 (Fig 3C). The loop and helix residues 337-357, which are disordered in all apo-TDO2 structures, are clearly visible in the holo-TDO2 structure (Fig 3C). This is in keeping with strong stabilizing interactions between hTDO2 and heme, including multiple hydrophobic interactions and the binding of His328 to the heme iron. The binding of AMT to the apo-TDO2 exosite (not shown) matches the published binding mode of tryptophan to holo-TDO2 [19].

### Structure of the inhibitor Cpd-4 bound to apo-TDO

The 2.61 Å resolution structure of Cpd-4 in complex with apo-TDO2 reveals multiple binding interactions (Fig 4) that explain its 500-fold higher potency compared to Rac-1 (15 nM vs 7.5 µM, respectively, in the cellular TDO2 assay). Cpd-4 is a mixture of two diastereomers, of which only ethyl (*R*)-9-(3-(2-isopropylphenyl)-1-((*S*)-1-phenylethyl)ureido)-2-methoxy-4-oxo-6,7,8,9-tetrahydro-4*H*-pyrido[1,2-*a*]pyrimidine-3-carboxylate is bound in the co-crystal structure. Both benzyl rings of the inhibitor are bound in hydrophobic pockets formed by phenylalanine, leucine and tryptophan side chains. The benzyl rings are bridged by a central urea linker that forms hydrogen bonds with the side chains of His328 and His76. Both histidine residues also play important roles in the function of holo-TDO2, with His328 anchoring the heme via its iron atom and His76 binding the substrate tryptophan. The backbone nitrogen of Ser155 forms an additional bifurcated hydrogen bond interaction with the inhibitor. The larger fused ring substituent binds between loop 145-155 and His328 and may serve to stabilize the loop conformation and shield the hydrogen bond between the inhibitor and His328 from competing water molecules.

**Fig 4.**
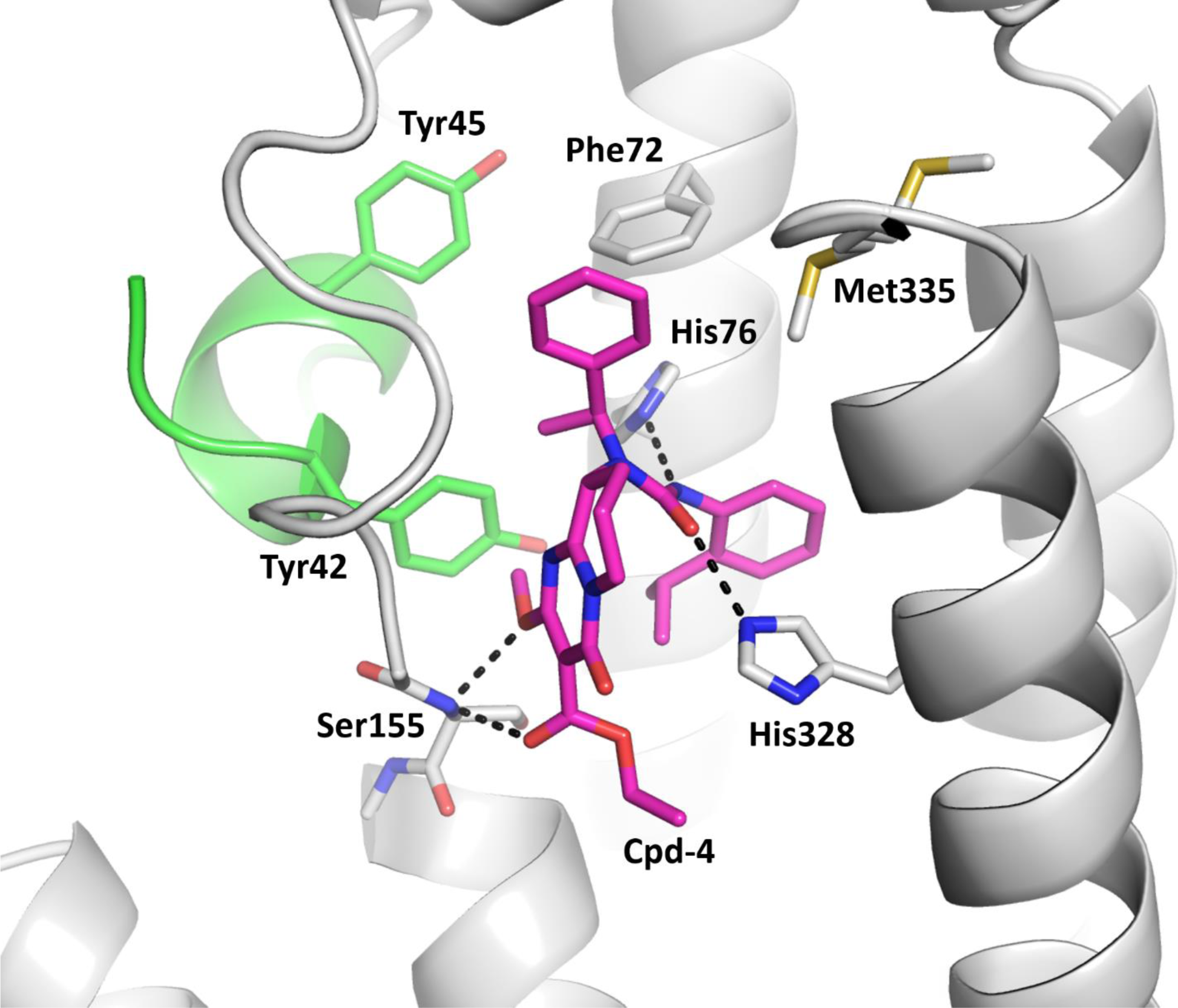
Binding interactions of the inhibitor Cpd-4 (magenta) in the heme binding pocket of apo-TDO. The binding pocket of TDO2 is formed by two TDO2 monomers, shown in grey and green cartoon. Hydrogen bond interactions between the inhibitor and TDO2 are shown as black dashes.

Rac-1 and Cpd-4 induce different conformations of the apo-TDO2 heme binding pocket (Figs 5 and 6). An overlay of both inhibitor complexes shows a significant rearrangement of loop residues 145-155. In uninhibited apo-TDO, loop 145-155 adopts an approximately intermediate conformation between the open conformation of Cpd-4 and the closed conformation of Rac-1 (S1 Fig 5).

**Fig 5.**
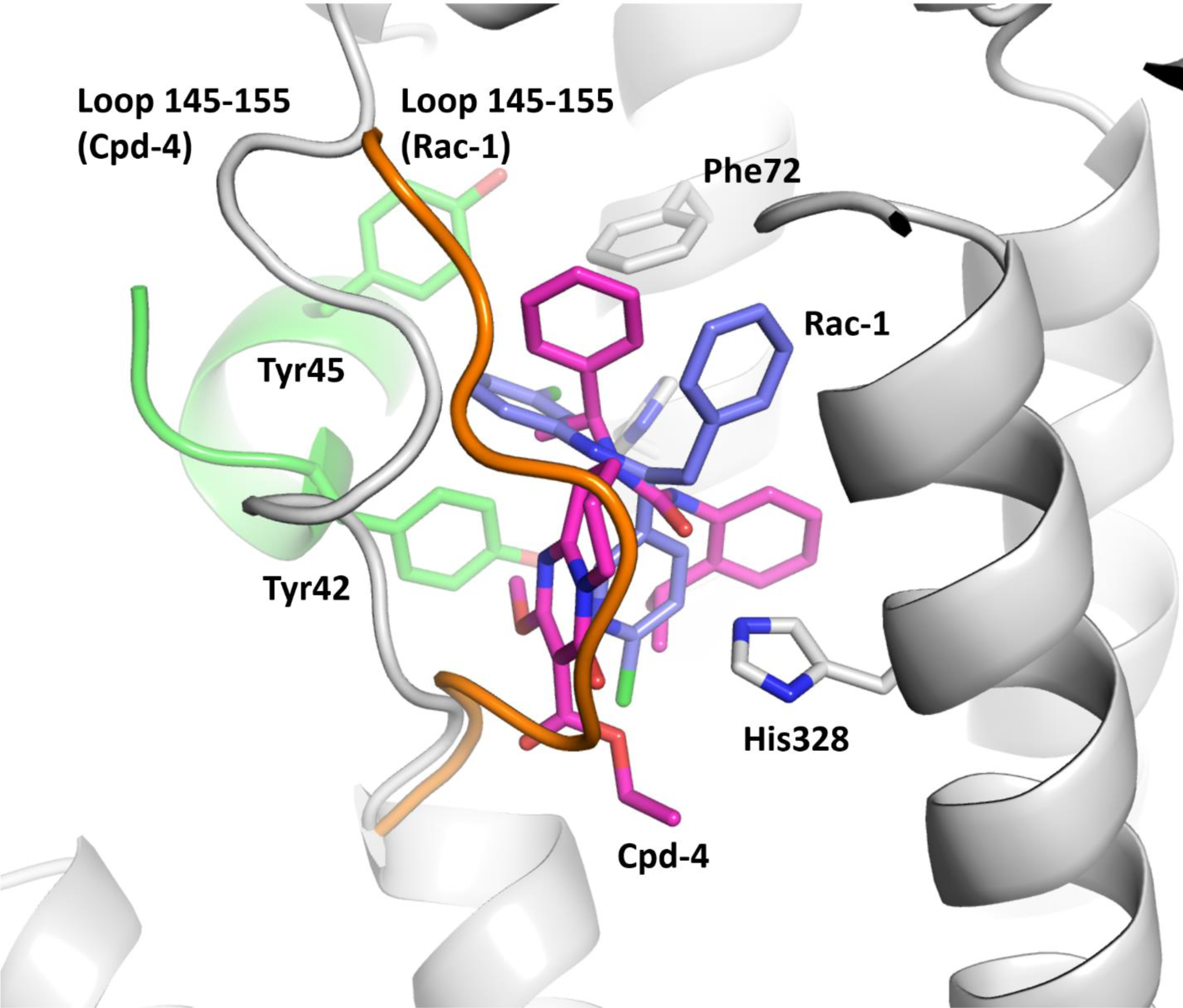
Superposition of the TDO2 inhibitor complexes of Rac-1 (cyan) and Cpd-4 (magenta) in the heme binding pocket of apo-TDO. Loop 145-155 of the Cpd-4 structure is shown in grey, with the more closed conformation of the loop in the Rac-1 structure shown in orange. The binding pocket of TDO2 is formed by two TDO2 monomers, shown in grey and green cartoon.

**Fig 6.**
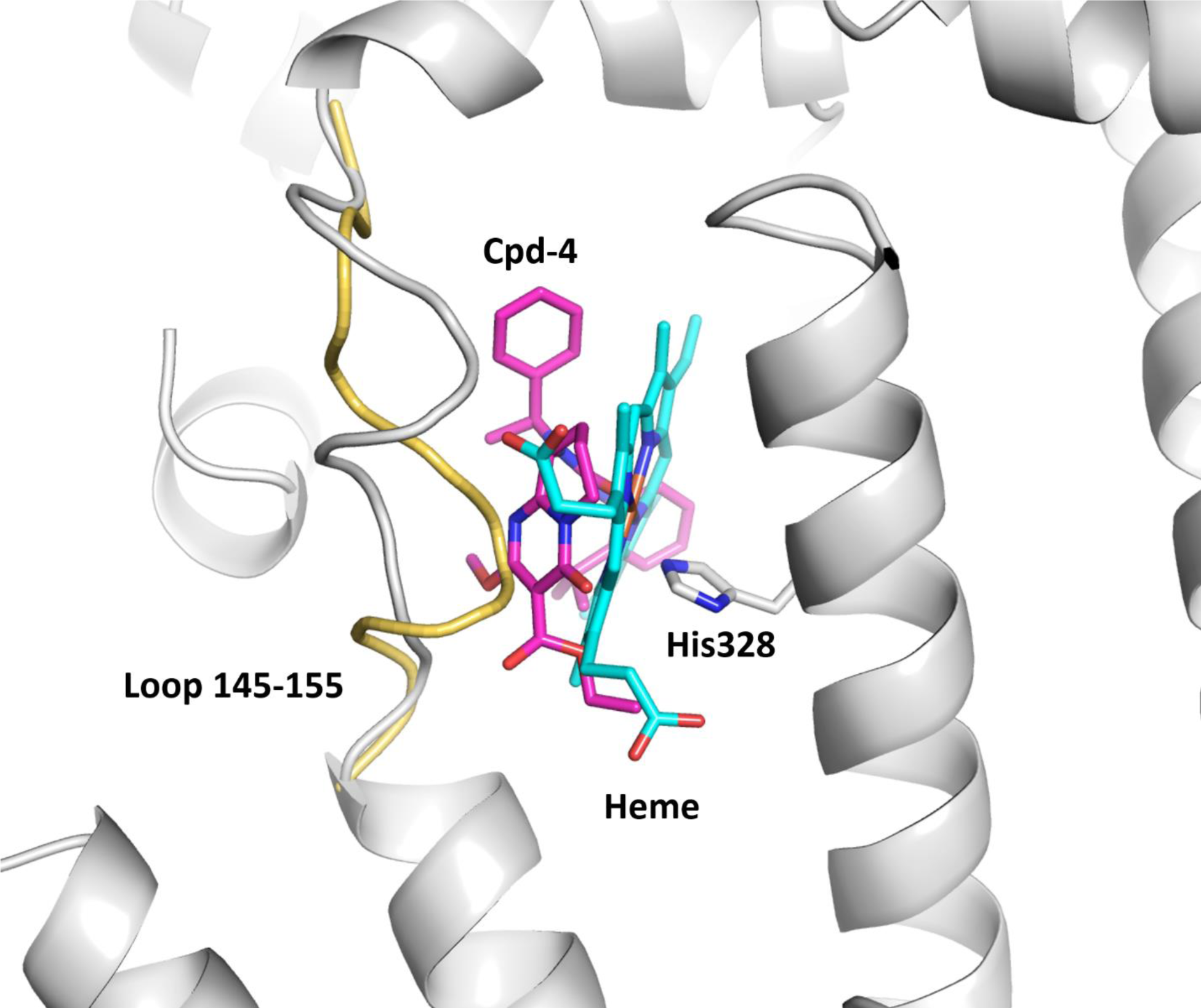
Superposition of the TDO2 inhibitor complexes of Cpd-4 (magenta) and heme (cyan) in the heme binding pocket of apo-TDO. Loop 145-155 of the Cpd-4 structure is shown in grey, with the loop of the holo-TDO2 structure shown in yellow.

To allow a comparison of the structures of apo-TDO2 inhibitor complexes with uninhibited structures, all with AMT bound in the exosite, the structure of AMT bound to uninhibited apo-TDO2 has also been solved (PDB 8QV7, space group C2). This structure is very similar to the published structure of apo-TDO2 without AMT (PDB entry ID 4PW8, space group P2_1_2_1_2_1_) [20] apart from a small conformational shift of residues 148-152.

## Discussion

We describe, for the first time to our knowledge, inhibitors binding to the apo form of TDO2 and inhibiting cellular TDO2 activity, together with the co-crystal structures of the inhibitors bound to apo-TDO2. While a former virtual screening to identify novel IDO1/TDO2 dual inhibitor scaffolds indicated that one of the hits might preferentially bind to the apo enzyme as revealed by molecular docking, experimental evidence to support the finding was lacking [21]. We believe that targeting the apo form of TDO2 offers a promising alternative approach for the development of therapeutic TDO2 inhibitors. Targeted inhibition of the apo form has been applied successfully to the structurally unrelated heme enzyme IDO1 [16], which together with TDO2 catalyzes the rate limiting first step of the kynurenine pathway.

Rational holo-TDO2 inhibitor development is hampered by a poor correlation between biochemical and cellular assay potencies, with inhibitors reported thus far showing limited cellular potency [8]. As demonstrated by the high potency of Cpd-4, targeting the apo form of the enzyme offers the possibility to generate more potent TDO2 inhibitors both as tools to study the kynurenine pathway and as potential therapeutics. Instead of competing with the Trp substrate, apo-TDO2 inhibitors compete with the insertion of the heme cofactor itself. The mechanism, timing, and reversibility of heme insertion into TDO2 remain poorly understood and require further investigation [1, 22]. Heme lability has been proposed as a posttranslational regulatory mechanism of both IDO1 and TDO2 [1]. Labile heme is present in the cells at very low concentrations (nM to single-digit µM concentrations) [23] while intracellular Trp concentrations have been reported in the range of 0.1-0.6 mM [24, 25] offering a competitive advantage to apo as compared to holo-binders. Another potential benefit of inhibitors targeting apo-TDO2 is stabilization of the apo form, unlike holo-TDO2 inhibitors which likely shift the balance of cellular TDO2 towards the holo form. Following elimination of an apo-TDO2 inhibitor, we expect a longer recovery period for cellular TDO2 activity when heme (re)binding is required, as described for IDO1.

Our work has several limitations which need to be addressed with further experiments. The kinetics of heme association with (or active insertion into) TDO2 and its dissociation rate in living cells needs to be further explored, in a similar way to the recent studies on cytochrome P450 enzyme maturation [26]. The balance of apo- vs. holo-TDO2 in different cells and disease states also requires further investigation. Most importantly, the development of optimized, potent apo-TDO2 inhibitors and investigation of their *in vivo* efficacy will allow us to further evaluate the apo form of TDO2 as a target for small molecule therapeutics.

## Materials and methods

### Production of human apo-TDO2

To produce truncated TDO2 as the apo form devoid of heme, recombinant protein expression was carried out using an *E. coli hemB* (porphobilinogen synthase) mutant strain unable to synthesize heme (RP523, *E. coli* Genetic Stock Center, Yale CGSC#7199). *E. coli* RP523 carries an uncharacterized permeability mutation that renders the bacteria heme-permeable, allowing aerobic propagation in supplementing growth media containing hemin [30 µg/ml] [27, 28].

DNA encoding a truncated version of human TDO2 (P48775.1, amino acid (aa) 39-389) with a C-terminal hexa-histidine tag was codon optimized for expression in *E. coli* and synthesized by GeneArt (Thermo Fisher Scientific). The TDO2 construct was provided as an insert in the pMA-T vector backbone with NdeI and BamHI as flanking restriction sites for subcloning into the pDSNdeI (AMP^r^) protein expression vector [29]. The aa 39-389 truncation was designed based on the clearly ordered region of the holo-TDO2 structure (PDB entry 5TIA) [19], with the aim of improving the expression yield and the likelihood of crystallization by removal of flexible termini.

The pDSNdeI derivative encoding the TDO2 construct was transformed into RP523 pRep4 (*lacI, Kan^r^*). RP523 pRep4 pDSNdeI::TDO2 aa39-389-His6 (NdeI, BamHI) was first grown aerobically (Luria-Bertani (LB) medium, ampicillin 100 µg/mL, kanamycin 25 µg/mL, 0.2% glucose, 2 µg/ml hemin, 37 °C, 225 rpm, Kühner Labtherm incubator) to prepare an inoculum containing sufficient cells for anaerobic expression. Inocula prepared under aerobic conditions (2 µg/mL hemin) were harvested and resuspended in medium without hemin to minimize transfer of residual hemin to anaerobic expression cultures.

The medium for anaerobic protein expression (6 x 1 liter LB, ampicillin 100 µg/mL, kanamycin 25 µg/mL, 0.2% glucose distributed to 4 screw cap flasks with magnetic stirrer) was equilibrated in an anaerobic glove box (Airlock, Coy Laboratory). The bacterial expression cultures were grown in the screw capped tightly closed flasks with moderate stirring. Protein expression was induced in the anaerobic glove box (100 µg/mL isopropyl ß-D-1-thiogalactopyranoside (IPTG)) at an optical density at 600 nm wavelength (OD_600_) of approximately 0.5. 50 µg/ml of Trp was added as additional supplement. Cultures were further grown overnight at 16 °C with moderate stirring.

Expressing cells were harvested by centrifugation using a Fiberlite F12-6×500 Lex Rotor with a Sorvall RC6+ centrifuge (20 min, 6000 revolutions per minute (rpm) (6371 *g*), 4 °C) and the bacterial cell pellets were stored at −70 °C.

### Purification of human apo-TDO2 proteins

Bacterial cell pellets of expression cultures (∼19.3 g) were thawed on ice and resuspended in lysis buffer (50 mM Tris-HCl pH 8.0, 400 mM NaCl, 1 mM TCEP, 5% glycerol, EDTA free completed protease inhibitor (Roche), 0.2 mg/ml lysozyme, 25 U/ml Benzonase nuclease, 10 mM MgCl_2_) for IMAC (immobilized metal (Ni) affinity chromatography). Cell breakage was carried out by high pressure homogenization (29008 pounds per square inch (psi), equal to 200 MPa) using a fluidizer (Microfluidics MP110P with DIXC H10Z chamber). Bacterial cell lysate was cleared by centrifugation (30 min at 16000 rpm (30392 *g*), Fiberlite F21-8×50y rotor) and applied to a 5 ml HisTrap Column (VWR, Cytiva) mounted on an Aekta Purifier 100 FPLC system.

Protein was eluted with a linear gradient of increasing imidazole (0-500 mM) concentration over 20 column volumes with buffer (50 mM Tris-HCl pH 8.0, 400 mM NaCl, 1 mM TCEP, 5% glycerol, 500 mM imidazole). Fractions (4 ml) were monitored for presence of TDO2 by SDS-PAGE (calculated molecular mass 42940.24 Da; theoretical pI 7.09, EXPASY protein parameter calculator https://web.expasy.org/protparam). Eluate fractions were analyzed by SDS-PAGE and immunoblot using an anti-histidine tag antibody (Monoclonal Mouse IgG1 histidine tag antibody; R&D Systems, cat. no. MAB050). Fractions containing TDO2 were combined and dialyzed (molecular weight cutoff 3.5 kDa, Spectrum^TM^, 132725) overnight to remove imidazole (50 mM Tris-HCl pH 8.0, 50 mM NaCl, 1 mM TCEP, 5% glycerol).

The protein was further purified by anion exchange chromatography (HiTrap Q, VWR 17-1154-01) with a linear gradient of increasing NaCl salt concentration. Eluate fractions containing TDO2 were combined and concentrated to a protein concentration of 2 mg/ml (Amicon, molecular weight cutoff 10 kDa). Buffer exchange of the concentrated TDO2 sample was carried out by size exclusion chromatography using a HiLoad 16/60 Superdex 200 column equilibrated with running buffer (50 mM HEPES-NaOH, pH 8.0, 200 mM NaCl, 2 mM TCEP, 10% glycerol).

Protein concentration was determined spectrophotometrically using the absorbance at 280 nm and the calculated extinction coefficient. The same methods were used to express and purify a longer construct (His6-aa19-406, with aa406 corresponding to the wild type C-terminal residue), which was used for DSF and other experiments.

### Production of human holo-TDO2

Human holo-TDO2 was produced in a similar way to the previously described procedure [19]. Briefly, a construct encoding amino acids 19-406 of human TDO with an N-terminal six histidine tag linked by a PreScission protease cleavage site was expressed in *E. coli* BL21(DE3) cells. Following cell disruption and centrifugation, the supernatant was purified by nickel affinity chromatography (HisTrap HP), followed by buffer exchange and anion exchange purification (HiTrap Q). The pooled eluate fractions were concentrated and purified by size exclusion chromatography (HiLoad 16/60 Superdex 200). All chromatography steps were carried out using an Äkta Purifier. The purified protein in a final buffer of 50 mM sodium phosphate pH 8.0, 300 mM NaCl, 5% glycerol was stored at −80 °C for use in enzymatic assays.

### Differential scanning fluorimetry

Protein thermal stability and small molecule ligand binding of TDO2 *in vitro* was studied using differential scanning fluorimetry (DSF, also known as thermal shift assay)[30]. Assay ingredients and ligands were combined to obtain a mixture composed of 10 µM TDO2 and 10x SYPRO Orange, in assay buffer (50 mM HEPES-NaOH pH 8.0, 200 mM NaCl, 2 mM TCEP, 10% glycerol) and 2% final DMSO concentration. Small molecule chemical ligands were taken from DMSO stock solution (100 mM, 10 mM) and diluted in DMSO transferred to wells of PCR Plates (Micro Amp Fast 96-well Reaction Plates, Applied Biosystems) to ensure matching final DMSO concentrations.

DSF was carried out with a StepOnePlus^TM^ real time PCR system. Thermal denaturation of TDO2 was monitored with a ramp rate of 1 K/min. Protein melting curves were evaluated using the Thermal Shift Software v 1.3. (Thermo Fisher Scientific) to derive differences in midpoint melting-temperatures (derivative model). The thermal shift (ΔTm) is documented as the difference between the reference midpoint melting temperature of proteins (containing 500 µM AMT) in the presence of the ligand vehicle (DMSO) and in the presence of ligand.

### Holo-TDO2 UV-VIS absorbance

A UV−visible spectra study was performed to evaluate the binding of the compounds. The holo-TDO2 absorbance spectra were measured using a BioTek Synergy multimode reader (Agilent) for wavelengths from 350 to 600 nm in 1 nm increments. 5 μM holo-TDO2 was incubated at 37 °C with 20 μM compounds or DMSO in phosphate buffer supplemented with 0.01% Tween-20 in a 96 well UV compatible plate (Greiner PS microplate #781101) and absorbance was measured before (t t=0h) and after 2-6 h of incubation.

### Crystallization and structure determination

All crystals were grown using the sitting drop vapour diffusion method at 293 K. The addition of AMT to apo-TDO2 was essential to obtain high quality crystals.

#### Apo-TDO2 with AMT

Truncated apo-TDO2 at a concentration of 20.5 mg/ml (480 µM) containing 2 mM AMT was crystallized using 50 mM HEPES-NaOH pH 8.0, 200 mM NaCl, 2 mM TCEP and 10% glycerol as a reservoir solution.

#### Apo-TDO2 with AMT and Rac-1

Truncated apo-TDO2 at a concentration of 5.3 mg/ml (124 µM) containing 2 mM AMT, was crystallized after incubation with 1 mM inhibitor from a 10 mM DMSO stock for two hours. The reservoir solution contained 12.5% (w/v) polyethylene glycol (PEG) 4000, 20% (v/v) 1,2,6-hexanetriol, 20 mM each of arginine, threonine, histidine, 5-hydroxylysine, and trans-4-hydroxy-l-proline, 100 mM glycylglycine/2-amino-2-methyl-1,3-propanediol (AMPD) pH 8.5 (condition G10 of the Morpheus II Screen)[31].

#### Apo-TDO2 with AMT and Cpd-1

Truncated apo-TDO2 at a concentration of 5.3 mg/ml (124 µM) containing 2 mM AMT, was crystallized after incubation with 1 mM inhibitor from a 10 mM DMSO stock for two hours. The reservoir solution contained 15% (w/v) PEG 3K, 20% (v/v) 1,2,4-butanetriol, 1% (w/v) nondetergent sulfobetaine (NDSB) 256, 20 mM each of xylitol, D-fructose, D-sorbitol, myo-inositol, and L-rhamnose monohydrate, 0.1 M N,N-bis(2-hydroxyethyl)-2-aminoethanesulfonic acid (BES)/triethanolamine (TEA) pH 7.5 (condition F5 of the Morpheus II Screen)[31].

#### Apo-TDO2 with AMT and Cpd-4

Truncated apo-TDO2 at a concentration of 5.3 mg/ml (124 µM) containing 2 mM AMT, was crystallized after incubation with 1 mM inhibitor from a 10 mM DMSO stock for two hours. The reservoir solution contained 10% (w/v) PEG 4000, 20% (v/v) glycerol, 0.03M each of sodium nitrate, disodium hydrogen phosphate, ammonium sulfate, 0.1M MES/imidazole pH 6.5 (condition C3 of the Morpheus I Screen)[31].

#### X-ray data collection and crystal structure determination

All crystals were harvested in loops and cryocooled directly in liquid nitrogen. Data collection was carried out at beamline X06DA/PXIII of the Swiss Light Source, Villigen, Switzerland (uninhibited TDO2 and the Rac-1 and Cpd-4 complexes) and at beamline ID23-1 of the ESRF, Grenoble, France (Cpd-1 complex). X-ray diffraction data were processed using autoPROC [32]. The structures were solved using Dimple and Phaser [33] for automated molecular replacement, Coot [34] for rebuilding and superpositioning, and BUSTER [35] for structure refinement and calculation of ligand RSCCs. Final 2mFo-DFc electron density maps were used for RSCC calculation, where m is the figure of merit and D is the sigma-A weighting factor. Ligand constraint files were created using Grade2 [36]. Figures were created using PyMOL [37]. The data processing and refinement statistics are shown in Supplementary Table 1.

The structures were deposited in the Protein Data Bank (PDB) with entry IDs 8QV7 (AMT), 8R5Q (AMT and Rac-1), 8R5R (AMT and Cpd-1) and 9EZJ (AMT and Cpd-4). The raw data (diffraction images) used for all structures were deposited at proteindiffraction.org and are accessible via the PDB entry.

### Assays

#### TDO2 cellular activity assay

SW48 cells (ATCC, CCL-231) were used to measure the TDO2 inhibitory activity of compounds and were routinely maintained in DMEM high glucose/GlutaMAX^TM^/pyruvate (Gibco, 10569010) 90% (v/v), FCS (Gibco, 10500, heat inactivated)10% (v/v), Penicillin/streptomycin (Gibco, 15140) 1% (v/v). SW48 cells were seeded in 384 well plates at a density of 8000 cells in 45 µl per well. Plates were incubated at 37 °C / 5% CO_2_ for 24 hours.

On the next day, 10 µl compound in serial dilutions (tested concentration range 0.5 nM to 10 µM) and 200 µM Trp were added. After 24 hours of incubation at 37 °C / 5% CO_2_, 3 µl of the supernatant per well was transferred into 25 µl water per well in a 384 deep well plate.

The 384 deep well plate containing 3 µl supernatant and 25 µl H_2_O per well were further processed for LCMS. After the addition of 100 µl Trp-(indole-d5) (Sigma 615862) at 200 nM in methanol, the 384 deep well plates were centrifuged for 10 minutes at 3220 *g* at 4 °C, 75 µl H_2_O was added per well and plates centrifuged again for 10 minutes at 3220 *g* at 4 °C. NFK and kynurenine were quantified by LCMS, normalized to the internal standard Trp-(indole-d5) and the sum was calculated.

Samples with 0.2% DMSO (0% effect) and the reference holo-TDO2 inhibitor 680C91[38] (100% effect) were used as control samples to set the parameters for the non-linear regression necessary for the determination of the IC_50_ for each compound and the determination of the Z’ factor for quality control. For each compound concentration the percentage of activity compared to 0% and 100% effect was calculated as average ± standard deviation (each concentration measured in duplicate). IC_50_ values and curves were generated with XLfit software (IDBS) using Dose-Response One Site model 203 (4 Parameter Logistic Model. Parameter A locked to 0 and B to 100). A serial dilution of the TDO2 inhibitor was included on each plate of the experiment and its IC50 was calculated. In the reported experiments, the deviation factor of the IC_50_ value of the reference TDO2 inhibitor was <3 versus the actual arithmetic mean and Z′ was >0.5.

As inhibition of NFK and KYN production can simply be an effect of cytotoxicity, a viability assay (CellTiter-Glo 2.0 Luminescent Cell Viability Assay, Promega Catalog # G9243) was performed in parallel. CellTiter-Glo reagent was added (25 µl/well) to cell plates, incubated for 15 minutes at room temperature in the dark and luminescence was measured with the EnVision Multilabel Reader from Perkin Elmer according to the manufacturer’s instructions. The luminescent signal is proportional to the amount of ATP present. The amount of ATP is directly proportional to the number of viable cells present. Samples with 0.2% DMSO (0% effect) and a toxic reference compound (100% effect) were used as control samples to set the parameters for the non-linear regression. For each compound concentration the percentage of activity compared to 0% and 100% effect was calculated as average ± STDEV (each concentration measured in duplicate). Cytotoxicity IC_50_ values and curves were generated with XLfit software (IDBS) using Dose-Response One Site model 203 (4 Parameter Logistic Model with parameter A locked to 0 and B to 100).

#### Binding to apo-TDO2 in living cells using NanoBRET

The human TDO2 gene (gene bank NM_005651, CDS: 64-1284, encoding the protein TDO) was synthesized and cloned into the lentiviral expression plasmid pLV.EF1 (Flash Therapeutics, Toulouse, France). The NanoLuc coding sequence including linker was obtained from the pNLF1-C plasmid (Promega) and inserted at the C-terminus of the TDO2 gene sequence (Flash Therapeutics, Toulouse, France) in the pLV.EF1 plasmid. The presence of an IRES element in pLV.EF1 allowed the bicistronic expression of TDO2-NLuc and a puromycin resistance gene. Lentiviral particles were produced at Flash Therapeutics and used to transduce MCF7 cells (NCI/No. 0507288) which have been confirmed to not express TDO2. TDO2-NLuc transduced MCF7 cells were selected by puromycin (GIBCO A11138-02). Functional expression of TDO2 was assessed by Western Blot and monitoring L-tryptophan conversion to NFK and kynurenine (see TDO2 cellular activity assay). NLuc expression was tested by luminescence measurement at 470 nm on the Envision plate reader using the Nano-Glo Luciferase Assay (Promega).

To test binding of TDO2 apo-binders in living cells, an apo-TDO2 binding molecule (structure not disclosed) was coupled to the NanoBRET 590SE (Promega) dye to create a fluorescent tracer. The tracer was confirmed to inhibit TDO2 activity in the SW48 cellular assay (S1 Fig 2) and tested for binding to apo-TDO2 in MCF7 cells expressing TDO-NLuc by NanoBRET under equilibrium conditions. In brief, MCF7-TDO-NLuc cells were trypsinized, resuspended in OptiMEM medium (GIBCO 11058-021) without additives and seeded in non-binding surface 384 well plates (Greiner 78190). Serially diluted tracer and fixed concentrations of tracer corresponding to EC_50_ and EC_80_ were added to cells in the presence or absence of 10 µM competing compound and plates were equilibrated for 2 h at 37 °C before BRET measurements. To measure BRET, plates were placed for 15 min at RT to equilibrate before NanoBRET NanoGlo Substrate (Promega) was added, and filtered luminescence was measured on a BMG LABTECH Clariostar luminometer using the Monochromator settings of the donor channel 470 nm/40 and acceptor channel 625 nm/50 with 1 s integration time and gain settings of 3,600 each.

To generate raw BRET ratio values, the acceptor emission value was divided by the donor emission value for each sample. The raw BRET units were converted to milliBRET (mBRET) units by multiplying each raw BRET value by 1,000. To correct for background, the BRET ratio in the absence of tracer (average of no-tracer control samples) was subtracted from the BRET ratio of each sample.

Apparent K_D_ and tracer affinity values were determined using curve fits in GraphPad Prism with the equation (equation (1)): Y = B_max_ * X / (K_D_ + X), where B_max_ is the maximum specific binding achieved (measured as BRET ratio), X is the tracer concentration and Y is the corresponding BRET ratio. In addition to K_D_ (EC_50_) determination, the equation was used to determine EC_80_ (0.8/0.2 * K_D_). EC_50_ and EC_80_ were determined in a pretest without competing compound and corresponded to tracer concentrations of 6 nM and 30 nM, respectively. Tracer titrations were run in each experiment and the GraphPad “Find ECanything” curve fitting was used to determine the actual EC corresponding to 6 nM and 30 nM Tracer concentration used in competition with compound.

## Acknowledgements

The authors would like to acknowledge Céline Potot, Marina Dos Santos, Thomas Lefebvre and François Le Goff for protein production, enzyme assays and mass spectrometry. The authors would also like to acknowledge the staff of the Swiss Light Source, Villigen, Switzerland, the ESRF, Grenoble, France, and Expose GmbH, Switzerland for their assistance in X-ray data collection.

## Abbreviations

AMPD: glycylglycine/2-amino-2-methyl-1,3-propanediol
AMT: alpha-methyl-L-tryptophan
BRET: bioluminescence resonance energy transfer
BSA: bovine serum albumin
DSF: differential scanning fluorimetry
IDO1: indoleamine-2,3-dioxygenase
KYN: kynurenine
LB: Luria-Bertani
NFK: N-formyl-kynurenine
NO: nitric oxide
psi: pounds per square inch
rpm: revolutions per minute
RSCC: real space correlation coefficient
TDO2: tryptophan-2,3-dioxygenase
Trp: L-tryptophan

## Supporting information captions

S1 Text. Protein Expression. Amino acid sequence of short (aa39-389) and long (aa19-406) TDO. Colour code: black, TDO2 sequence; blue, additional amino acids introduced as a linker or during subcloning; red, 6 histidine tag.

S1 Table. Data collection and refinement statistics for TDO2 bound to AMT alone, AMT and Rac-1, and AMT and Cpd-1.

S1 Figure. IC_50_ determination of inhibitor Rac-1 and Cpd-4 in the TDO2 SW48 cellular assay.

S2 Figure. IC_50_ determination of tracer in the TDO2 SW48 cellular assay.

S3 Figure. Fo-Fc omit electron density maps (green, 2.62 Å resolution) contoured at 3 sigma level, showing Rac-1 bound to TDO2 chains A-D (matching figure lettering). Rac-1 is shown as sticks with atom color magenta (C), blue (N) and green (Cl).

S4 Figure. Fo-Fc omit electron density maps (green, 3.08 Å resolution) contoured at 3 sigma level, showing Cpd-1 bound to TDO2 chains A-D (matching figure lettering). Cpd-1 is shown as sticks with atom color magenta (C), blue (N) and green (Cl).

S5 Figure. Fo-Fc omit electron density maps (green, 2.61 Å resolution) contoured at 3 sigma level, showing Cpd-4 bound to TDO2 chains A-D (matching figure lettering). Cpd-4 is shown as sticks with atom color magenta (C), blue (N) and red (O).

## References

1. Biswas P, Stuehr DJ. Indoleamine dioxygenase and tryptophan dioxygenase activities are regulated through control of cell heme allocation by nitric oxide. J Biol Chem. 2023;299(6): 104753.

2. Opitz CA, Somarribas Patterson LF, Mohapatra SR, Dewi DL, Sadik A, Platten M, et al. The therapeutic potential of targeting tryptophan catabolism in cancer. Br J Cancer. 2020;122(1): 30–44.

3. Ortiz-Meoz RF, Wang L, Matico R, Rutkowska-Klute A, De la Rosa M, Bedard S, et al. Characterization of Apo-Form Selective Inhibition of Indoleamine 2,3-Dioxygenase*. Chembiochem. 2021;22(3): 516–22.

4. Liu W, Zou Y, Li K, Zhong H, Yu L, Ge S, et al. Apo-Form Selective Inhibition of IDO for Tumor Immunotherapy. J Immunol. 2022;209(1): 180–91.

5. Crosignani S, Bingham P, Bottemanne P, Cannelle H, Cauwenberghs S, Cordonnier M, et al. Discovery of a Novel and Selective Indoleamine 2,3-Dioxygenase (IDO-1) Inhibitor 3-(5-Fluoro-1H-indol-3-yl)pyrrolidine-2,5-dione (EOS200271/PF-06840003) and Its Characterization as a Potential Clinical Candidate. J Med Chem. 2017;60(23): 9617–29.

6. Peng YH, Ueng SH, Tseng CT, Hung MS, Song JS, Wu JS, et al. Important Hydrogen Bond Networks in Indoleamine 2,3-Dioxygenase 1 (IDO1) Inhibitor Design Revealed by Crystal Structures of Imidazoleisoindole Derivatives with IDO1. J Med Chem. 2016;59(1): 282–93.

7. Lancellotti S, Novarese L, De Cristofaro R. Biochemical properties of indoleamine 2,3-dioxygenase: from structure to optimized design of inhibitors. Curr Med Chem. 2011;18(15): 2205–14.

8. Pei Z, Mendonca R, Gazzard L, Pastor R, Goon L, Gustafson A, et al. Aminoisoxazoles as Potent Inhibitors of Tryptophan 2,3-Dioxygenase 2 (TDO2). ACS Med Chem Lett. 2018;9(5): 417–21.

9. Zhang Y, Li Y, Chen X, Chen X, Chen C, Wang L, et al. Discovery of 1-(Hetero)aryl-beta-carboline Derivatives as IDO1/TDO Dual Inhibitors with Antidepressant Activity. J Med Chem. 2022;65(16): 11214–28.

10. Zhang Y, Hu Z, Zhang J, Ren C, Wang Y. Dual-target inhibitors of indoleamine 2, 3 dioxygenase 1 (Ido1): A promising direction in cancer immunotherapy. Eur J Med Chem. 2022;238: 114524.

11. Yang L, Chen Y, He J, Njoya EM, Chen J, Liu S, et al. 4,6-Substituted-1H-Indazoles as potent IDO1/TDO dual inhibitors. Bioorg Med Chem. 2019;27(6): 1087–98.

12. Peng X, Zhao Z, Liu L, Bai L, Tong R, Yang H, et al. Targeting Indoleamine Dioxygenase and Tryptophan Dioxygenase in Cancer Immunotherapy: Clinical Progress and Challenges. Drug Des Devel Ther. 2022;16: 2639–57.

13. Naing A, Eder JP, Piha-Paul SA, Gimmi C, Hussey E, Zhang S, et al. Preclinical investigations and a first-in-human phase I trial of M4112, the first dual inhibitor of indoleamine 2,3-dioxygenase 1 and tryptophan 2,3-dioxygenase 2, in patients with advanced solid tumors. J Immunother Cancer. 2020;8(2).

14. Long GV, Dummer R, Hamid O, Gajewski TF, Caglevic C, Dalle S, et al. Epacadostat plus pembrolizumab versus placebo plus pembrolizumab in patients with unresectable or metastatic melanoma (ECHO-301/KEYNOTE-252): a phase 3, randomised, double-blind study. Lancet Oncol. 2019;20(8): 1083–97.

15. Balog A, Lin TA, Maley D, Gullo-Brown J, Kandoussi EH, Zeng J, et al. Preclinical Characterization of Linrodostat Mesylate, a Novel, Potent, and Selective Oral Indoleamine 2,3-Dioxygenase 1 Inhibitor. Mol Cancer Ther. 2021;20(3): 467–76.

16. Nelp MT, Kates PA, Hunt JT, Newitt JA, Balog A, Maley D, et al. Immune-modulating enzyme indoleamine 2,3-dioxygenase is effectively inhibited by targeting its apo-form. Proc Natl Acad Sci U S A. 2018;115(13): 3249–54.

17. Seegers N, van Doornmalen AM, Uitdehaag JC, de Man J, Buijsman RC, Zaman GJ. High-throughput fluorescence-based screening assays for tryptophan-catabolizing enzymes. J Biomol Screen. 2014;19(9): 1266–74.

18. Robers MB, Dart ML, Woodroofe CC, Zimprich CA, Kirkland TA, Machleidt T, et al. Target engagement and drug residence time can be observed in living cells with BRET. Nat Commun. 2015;6: 10091.

19. Lewis-Ballester A, Forouhar F, Kim SM, Lew S, Wang Y, Karkashon S, et al. Molecular basis for catalysis and substrate-mediated cellular stabilization of human tryptophan 2,3-dioxygenase. Sci Rep. 2016;6: 35169.

20. Meng B, Wu D, Gu J, Ouyang S, Ding W, Liu ZJ. Structural and functional analyses of human tryptophan 2,3-dioxygenase. Proteins. 2014;82(11): 3210–6.

21. Sari S, Tomek P, Leung E, Reynisson J. Discovery and Characterisation of Dual Inhibitors of Tryptophan 2,3-Dioxygenase (TDO2) and Indoleamine 2,3-Dioxygenase 1 (IDO1) Using Virtual Screening. Molecules. 2019;24(23): 4346.

22. Biswas P, Dai Y, Stuehr DJ. Indoleamine dioxygenase and tryptophan dioxygenase activities are regulated through GAPDH- and Hsp90-dependent control of their heme levels. Free Radic Biol Med. 2022;180: 179–90.

23. Gallio AE, Fung SS, Cammack-Najera A, Hudson AJ, Raven EL. Understanding the Logistics for the Distribution of Heme in Cells. JACS Au. 2021;1(10): 1541–55.

24. Kaper T, Looger LL, Takanaga H, Platten M, Steinman L, Frommer WB. Nanosensor detection of an immunoregulatory tryptophan influx/kynurenine efflux cycle. PLoS Biol. 2007;5(10): e257.

25. Tao R, Wang K, Chen TL, Zhang XX, Cao JB, Zhao WQ, et al. A genetically encoded ratiometric indicator for tryptophan. Cell Discov. 2023;9(1): 106.

26. Islam S, Jayaram DT, Biswas P, Stuehr DJ. Functional Maturation of Cytochromes P4503A4 and 2D6 Relies on GAPDH- and Hsp90-Dependent Heme Allocation. Journal of Biological Chemistry.

27. Li JM, Umanoff H, Proenca R, Russell CS, Cosloy SD. Cloning of the Escherichia coli K-12 hemB gene. J Bacteriol. 1988;170(2): 1021–5.

28. Woodward JJ, Martin NI, Marletta MA. An Escherichia coli expression-based method for heme substitution. Nat Methods. 2007;4(1): 43–5.

29. Stüber D, Matile H, Garotta G. 8 - System for High-Level Production in Escherichia coli and Rapid Purification of Recombinant Proteins: Application to Epitope Mapping, Preparation of Antibodies, and Structure—Function Analysis. In: Ivan L, Benvenuto P, editors. Immunological Methods: Academic Press; 1990. p. 121–52.

30. Niesen FH, Berglund H, Vedadi M. The use of differential scanning fluorimetry to detect ligand interactions that promote protein stability. Nat Protoc. 2007;2(9): 2212–21.

31. Gorrec F. The MORPHEUS II protein crystallization screen. Acta Crystallogr F Struct Biol Commun. 2015;71(Pt 7): 831–7.

32. Vonrhein C, Flensburg C, Keller P, Sharff A, Smart O, Paciorek W, et al. Data processing and analysis with the autoPROC toolbox. Acta Crystallogr D Biol Crystallogr. 2011;67(Pt 4): 293–302.

33. Collaborative Computational Project N. The CCP4 suite: programs for protein crystallography. Acta Crystallogr D Biol Crystallogr. 1994;50(Pt 5): 760–3.

34. Emsley P, Lohkamp B, Scott WG, Cowtan K. Features and development of Coot. Acta Crystallogr D Biol Crystallogr. 2010;66(Pt 4): 486–501.

35. Bricogne G, Blanc E, Brandl M, Flensburg C, Keller P, Paciorek W, et al. BUSTER. 2.11.8 ed. Cambridge, United Kingdom: Global Phasing Limited; 2017.

36. Smart OS, Womack TO, Sharff A, Flensburg C, Keller P, Paciorek W, et al. Grade2, version 1.4.0 Cambridge, United Kingdom: Global Phasing Ltd.; 2011 [Available from: https://www.globalphasing.com.

37. Schrodinger, LLC. The PyMOL Molecular Graphics System, Version 1.8. 2015.

38. Dolusic E, Larrieu P, Moineaux L, Stroobant V, Pilotte L, Colau D, et al. Tryptophan 2,3-dioxygenase (TDO) inhibitors. 3-(2-(pyridyl)ethenyl)indoles as potential anticancer immunomodulators. J Med Chem. 2011;54(15): 5320–34.

