## Supplementary Information for "Discovery and binding mode of small molecule inhibitors of the apo form of human TDO2"

#### Protein Expression

aa39-389

```
MGLIYGNYLH LEKVLNAQEL QSETKGNKIH DEHLFIITHQ AYELWFKQIL WELDSVREIF
QNGHVRDERN MLKVVSRMHR VSVILKLLVQ QFSILETMTA LDFNDFREYL SPASGFQSLQ
FRLLENKIGV LQNMRVPYNR RHYRDNFKGE ENELLLKSEQ EKTLLELVEA WLERTPGLEP
HGFNFWGKLE KNITRGLEEE FIRIQAKEES EEKEEQVAEF QKQKEVLLSL FDEKRHEHLL
SKGERRLSYR ALQGALMIYF YREEPRFQVP FQLLTSLMDI DSLMTKWRYN HVCMVHRMLG
SKAGTGGSSG YHYLRSTVSD RYKVFVDLFN LSTYLIPRHW IPKMNPTIHK FLEHHHHHH
```

aa19-406

```
MRGSHHHHHH GIDHMPVEGS EEDKSQTGVN RASKGGLIYG NYLHLEKVLN AQELQSETKG
NKIHDEHLFI ITHQAYELWF KQILWELDSV REIFQNGHVR DERNMLKVVS RMHRVSVILK
LLVQQFSILE TMTALDFNDF REYLSPASGF QSLQFRLLEN KIGVLQNMRV PYNRRHYRDN
FKGEENELLL KSEQEKTLLE LVEAWLERTP GLEPHGFNFW GKLEKNITRG LEEEFIRIQA
KEESEEKEEQ VAEFQKQKEV LLSLFDEKRH EHLLSKGERR LSYRALQGAL MIYFYREEPR
FQVPFQLLTS LMDIDSLMTK WRYNHVCMVH RMLGSKAGTG GSSGYHYLRS TVSDRYKVFV
DLFNLSTYLI PRHWIPKMNP TIHKFLYTAE YCDSSYFSSD ESD
```

### X-ray data processing and refinement statistics

| | | **AMT** | **Rac-1 / AMT** | **Cpd-1 / AMT** | **Cpd-4 / AMT** |
|---|---|---|---|---|---|
| **Data collection** | | | | | |
| Source | | SLS X06DA | SLS X10SA | ESRF ID23-1 | SLS X06DA |
| Space Group | | C2 | $P2_12_12_1$ | $P2_12_12_1$ | $P2_12_12$ |
| Wavelength (Å) | | 1.00 | 1.00 | 0.89 | 1.00 |
| Cell dimensions | a, b, c (Å)<br>α, β, γ (°) | 181.3, 90.8, 132.8<br>90.0, 120.8, 90.0 | 91.3, 132.5, 135.6<br>90.0, 90.0, 90.0 | 91.7, 132.5, 137.4<br>90.0, 90.0, 90.0 | 133.8, 156.2, 90.7<br>90.0, 90.0, 90.0 |
| Observed reflections | | 109032 | 269737 | 272168 | 291634 |
| Unique reflections | | 31931 | 39729 | 27716 | 43550 |
| Resolution (highest shell) (Å) | | 114.1-2.66 (3.05-2.66) | 94.8-2.62 (2.86-2.62) | 76.31-3.08 (3.24-3.08) | 45.33-2.61 (2.87-2.61) |
| Rpim (%) | | 12.2 (35.4) | 10.1 (145.1) | 5.6 (64.8) | 11.2 (57.5) |
| Rmerge (%) | | 19.1 (55.9) | 9.4 (133.3) | 17.1 (180.1) | 26.0 (126.8) |
| Mean I/σ(I) | | 3.2 (1.6) | 14.5 (1.5) | 9.8 (1.3) | 6.0 (1.5) |
| Completeness (%) | Spherical | 59.8 (9.0) | 74.8 (17.2) | 87.0 (31.0) | 74.7 (15.5) |
| | Ellipsoidal | 90.7 (51.5) | 94.8 (58.3) | 95.2 (60.6) | 94.2 (61.7) |
| Redundancy | | 3.4 (3.4) | 6.8 (6.4) | 9.8 (8.6) | 6.7 (6.7) |
| $CC_{0.5}$ | | (0.73) | (0.61) | (0.54) | (0.69) |
| **Refinement** | | | | | |
| Resolution (highest shell) (Å) | | 34.46-2.66 (2.86-2.66) | 75.75-2.62 (2.77-2.62) | 50.83-3.08 (3.18-3.08) | 45.33-2.61 (2.87-2.61) |
| $R_{work}$ (highest shell) (%) | | 22.9 (26.9) | 25.4 (34.0) | 22.8 (32.7) | 30.0 (37.4) |
| $R_{free}$ (highest shell) (%) | | 6.0 (36.5) | 26.7 (41.0) | 25.2 (32.5) | 33.0 (48.2) |
| B-factors (Å$^2$) | Protein | 34.5 | 79.0 | 83.7 | 38.7 |
| | Ligands | 27.8 | 76.6 | 82.3 | 33.4 |
| | Water | 13.2 | 52.3 | 53.8 | 8.9 |
| Rms deviation | Bond lengths (Å) | 0.007 | 0.007 | 0.007 | 0.007 |
| | Bond angles (°) | 0.83 | 0.83 | 0.82 | 0.81 |
| Ramachandran plot (%) | Favoured | 96.4 | 96.1 | 95.9 | 96.8 |
| | Allowed | 3.4 | 3.0 | 3.3 | 2.5 |
| | Disallowed | 0.2 | 0.9 | 0.8 | 0.7 |
| PBD entry ID | | 8QV7 | 8R5Q | 8R5R | 9EZJ |

Supplementary Table 1. Data collection and refinement statistics for TDO2, Rac-1 (racemate), Cpd-1 (single enantiomer of Rac-1) and Cpd-4. All structures contain AMT bound at the exosite of TDO2.

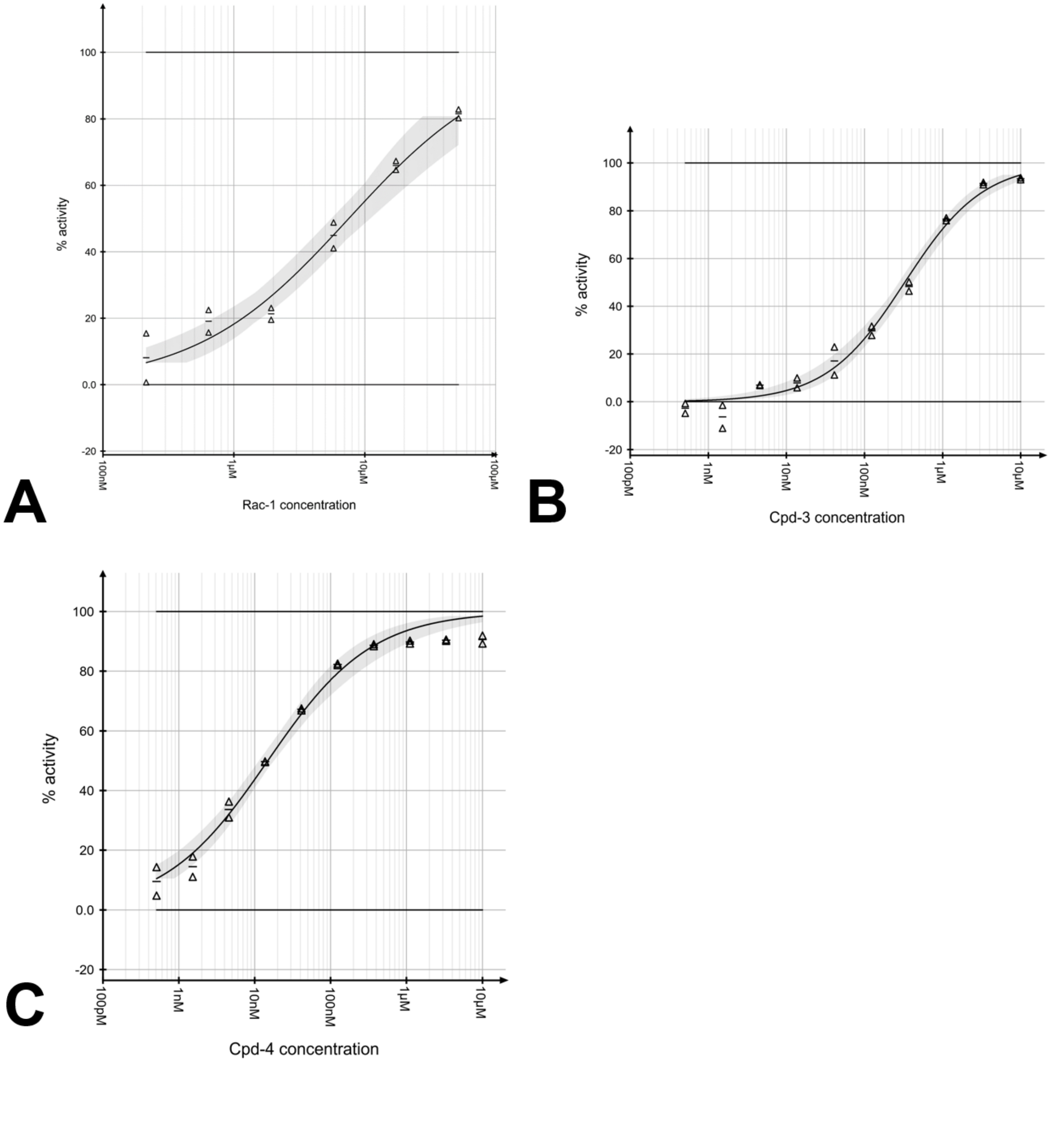


S1 Figure. $IC_{50}$ determination of inhibitor Rac-1 (A), Cpd-3 (B), and Cpd-4 (C) in the TDO2 SW48 cellular assay.

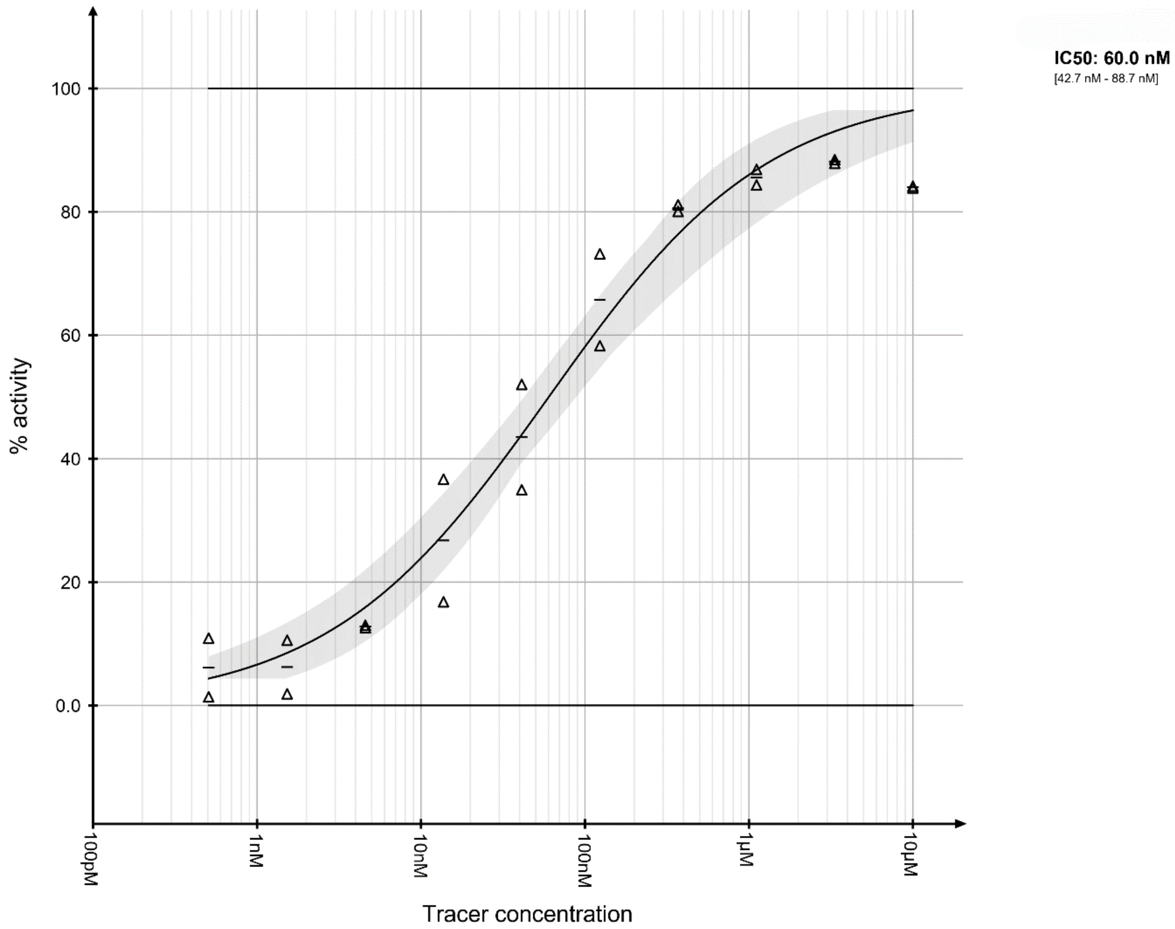

S2 Figure. $IC_{50}$ determination of tracer in the TDO2 SW48 cellular assay.

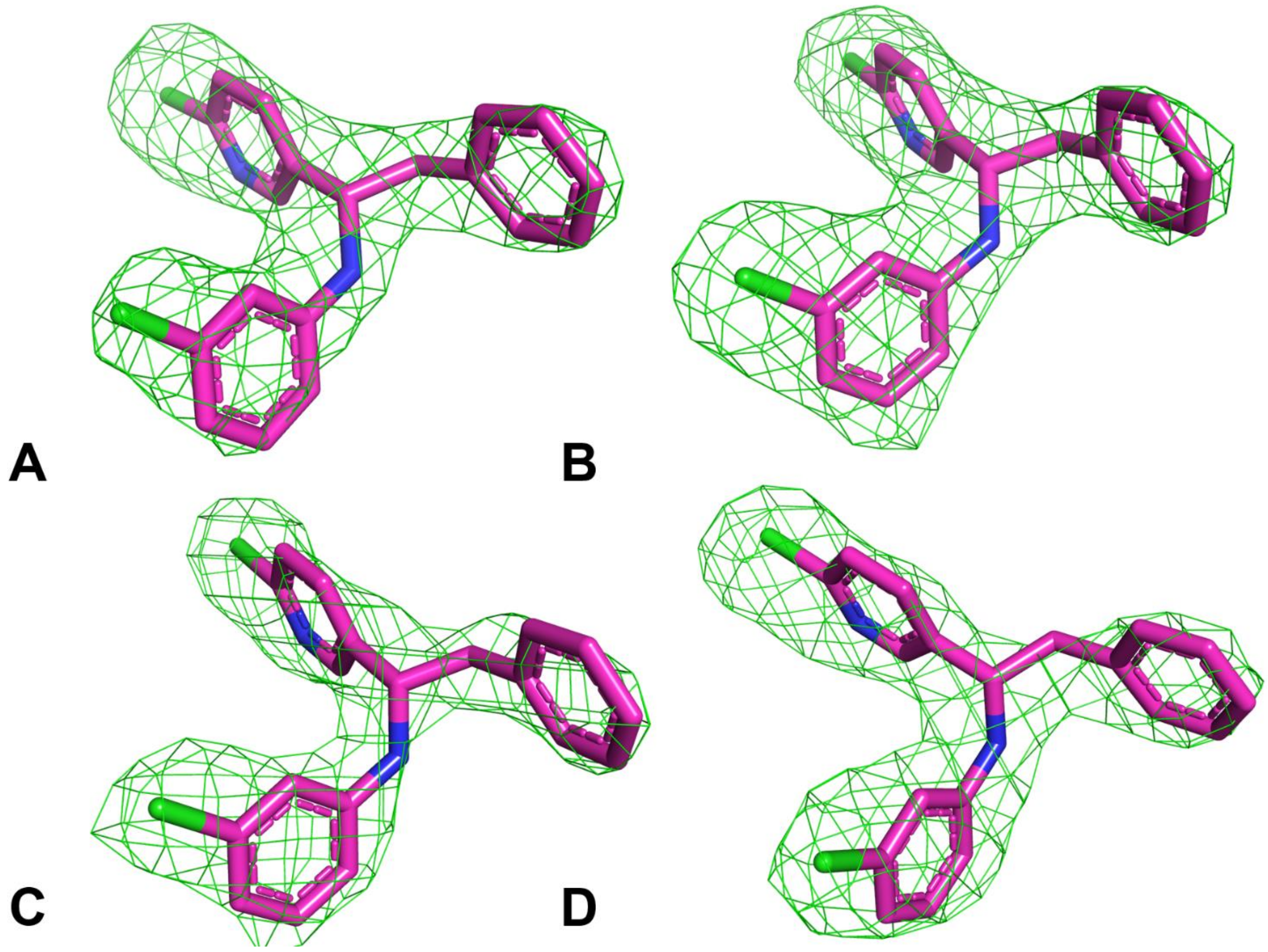


S3 Figure. Fo-Fc omit electron density maps (green, 2.62 Å resolution) contoured at 3 sigma level, showing Rac-1 bound to TDO2 chains A-D (matching figure lettering). Rac-1 is shown as sticks with atom color magenta (C), blue (N) and green (Cl).

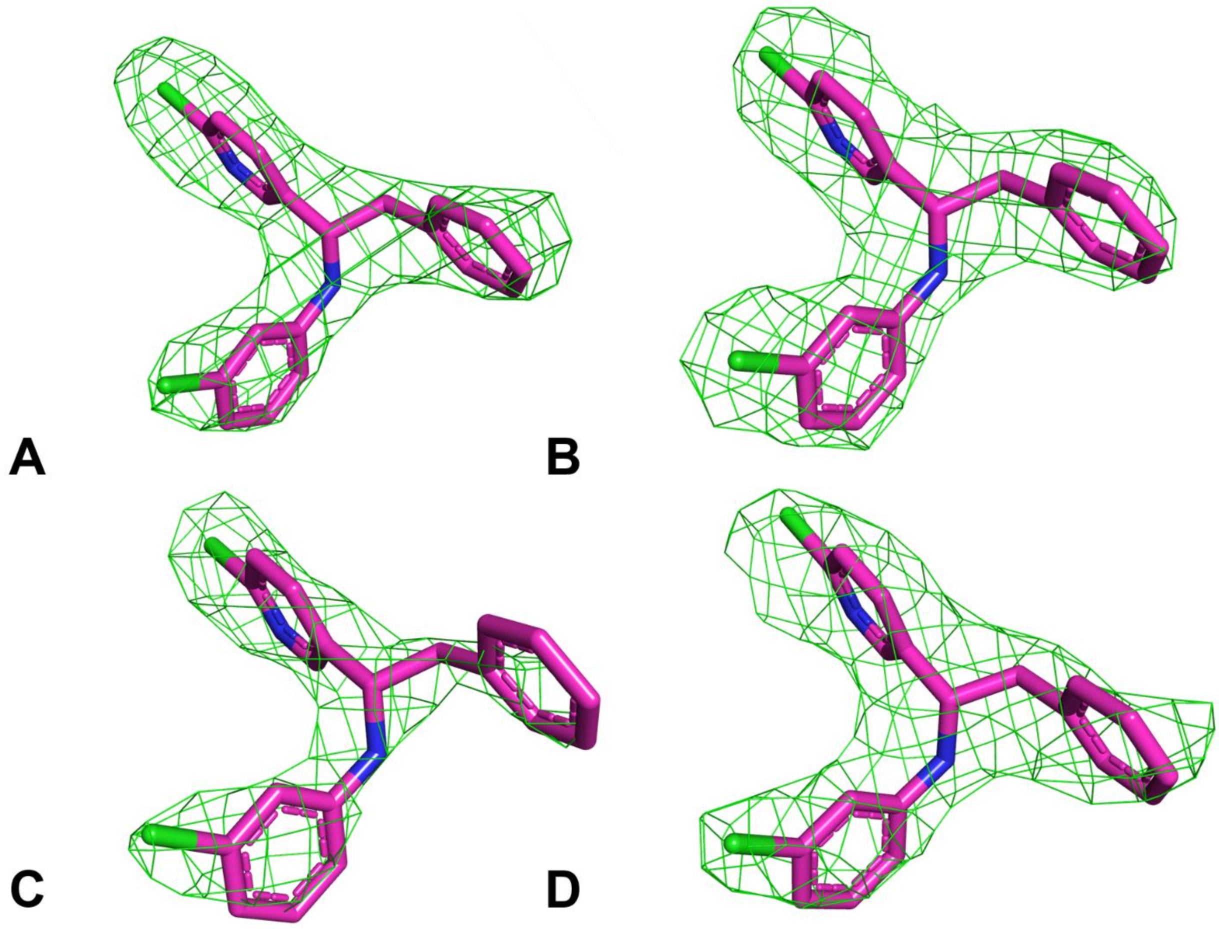


S4 Figure. Fo-Fc omit electron density maps (green, 3.08 Å resolution) contoured at 3 sigma level, showing Cpd-1 bound to TDO2 chains A-D (matching figure lettering). Cpd-1 is shown as sticks with atom color magenta (C), blue (N) and green (Cl).

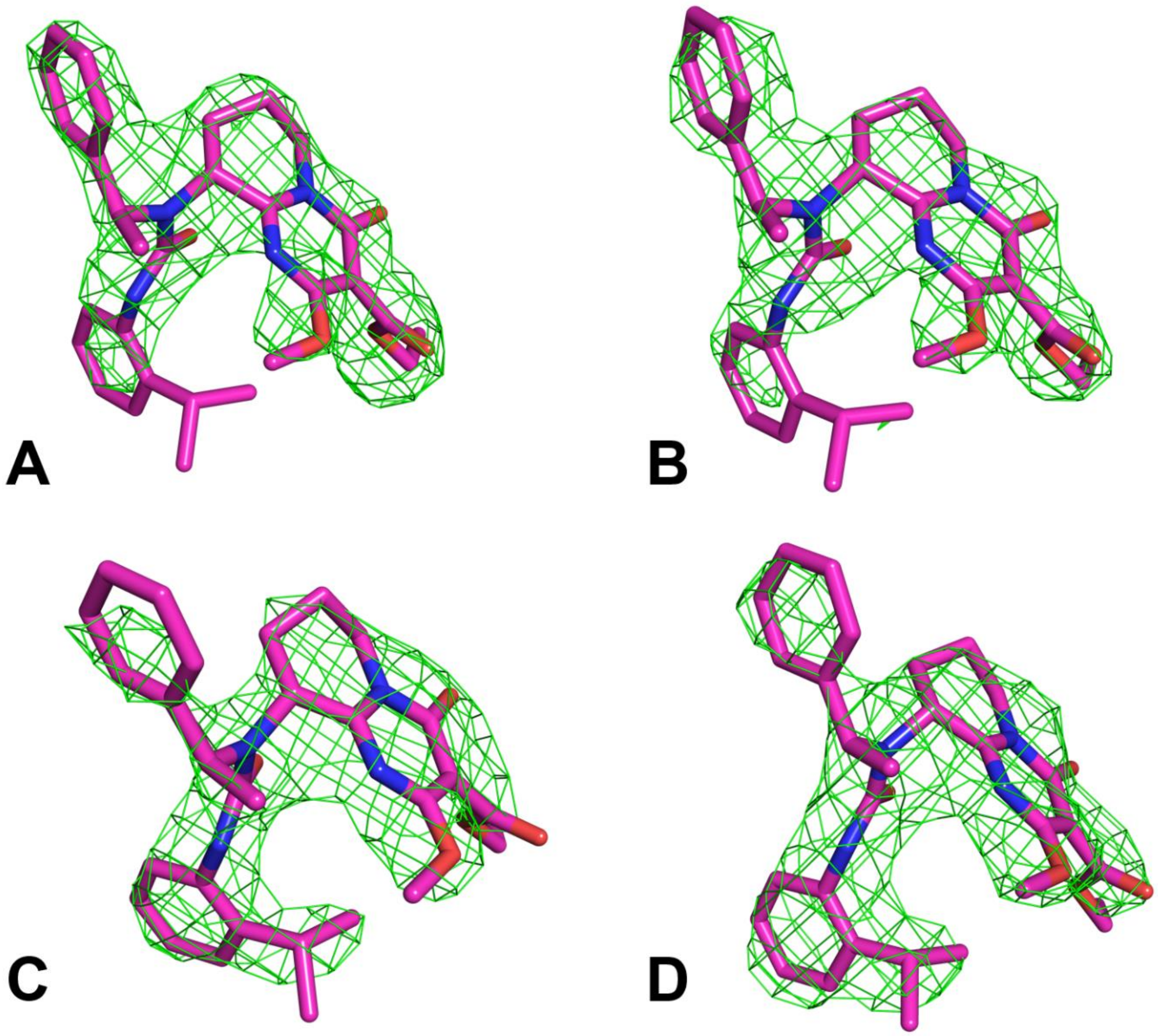


S5 Figure. Fo-Fc omit electron density maps (green, 2.61 Å resolution) contoured at 3 sigma level, showing Cpd-4 bound to TDO2 chains A-D (matching figure lettering). Cpd-4 is shown as sticks with atom color magenta (C), blue (N) and red (O).

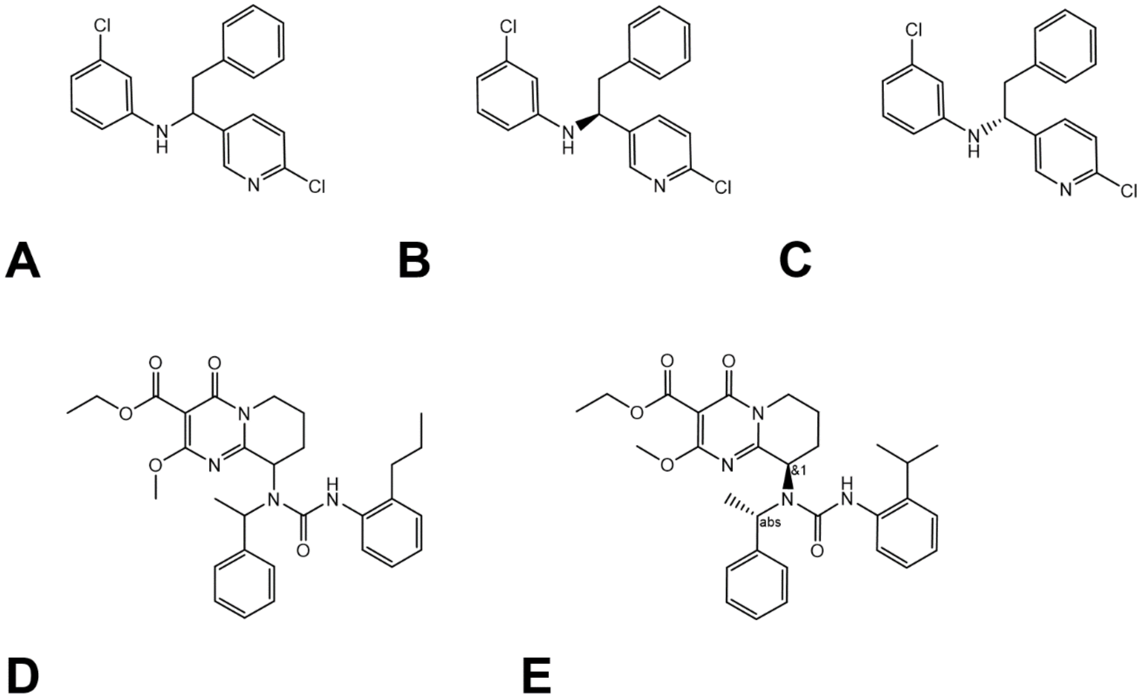


S6 Figure. Chemical structure of the racemate Rac-1 (A), its purified enantiomers Cpd-1 (B) and Cpd-2 (C), and the diastereomeric mixtures Cpd-3 (D) and Cpd-4 (E). The absolute configuration of Cpd-1 was tentatively assigned based on the crystal structure in complex with TDO2. Only the (R,S) diastereomer of Cpd-4 (as drawn) was observed in the TDO2 co-crystal structure.
